# Evidence for an RNAi-independent role of DICER-LIKE2 in conferring growth inhibition and basal antiviral resistance

**DOI:** 10.1101/2023.01.10.523401

**Authors:** Carsten Poul Skou Nielsen, Laura Arribas-Hernández, Lijuan Han, Stig Uggerhøj Andersen, Nathan Pumplin, Peter Brodersen

## Abstract

Higher plants encode four DICER-LIKE (DCL) enzymes responsible for the production of small non-coding RNAs which function in RNA interference (RNAi). Different RNAi pathways in plants effect transposon silencing, antiviral defense and endogenous gene regulation. *DCL2* acts genetically redundantly with *DCL4* to confer basal antiviral defense, but in other settings, *DCL2* has the opposite function of *DCL4*, at least in formal genetic terms. For example, knockout of *DCL4* causes growth defects that are suppressed by inactivation of *DCL2*. Current models maintain that the biochemical basis of both of these effects is RNAi via DCL2-dependent small interfering RNAs (siRNAs). Here, we report that neither DCL2-mediated antiviral resistance nor growth defects can be explained by silencing effects of DCL2-dependent siRNAs. Both functions are defective in genetic backgrounds that maintain high levels of DCL2-dependent siRNAs, either through specific point mutations in DCL2 or simply by reducing DCL2 dosage in plants heterozygous for *dcl2* knockout alleles. Intriguingly, however, all functions of DCL2 depend on it having some level of catalytic activity. We discuss this requirement for catalytic activity, but not for the resulting siRNAs, in the light of recent findings that reveal a function of DCL2 in activation of innate immunity in response to cytoplasmic double-stranded RNA.

## INTRODUCTION

Small non-coding RNAs are central players in the control of genetic information in eukaryotic organisms. Plants make pervasive use of small interfering RNAs (siRNAs) to defend against transposable elements and viruses (Ding and Voinnet, 2007; Law and Jacobsen, 2010), and of both siRNAs and microRNAs (miRNAs) to regulate endogenous gene expression programs during development and in response to changes in the environment (D’Ario et al., 2017; Song et al., 2019). Both classes of regulatory small RNA are produced from longer double-stranded (ds) precursor RNA molecules by multidomain DICER-LIKE (DCL) ribonucleases (Fukudome and Fukuhara, 2017). DCLs use two RNaseIII-domains for dsRNA cleavage and a Piwi-Argonaute-Zwille (PAZ) domain to help anchor the extremity of staggered dsRNA substrates and position the catalytic center in the correct distance from the dsRNA end to produce small RNA duplexes of a well-defined size between 21 and 24 nt depending on the exact DCL enzyme (Macrae et al., 2006; MacRae et al., 2007). In addition, all DCLs contain conserved DExH-box helicase domains (Song and Rossi, 2017) that may facilitate binding of blunt end substrates and processive action of the enzyme on long dsRNA substrates (Niladri et al., 2018). The staggered duplex product released upon DCL-mediated processing subsequently associates with an ARGONAUTE (AGO) protein to form a mature RNA Induced Silencing Complex (RISC) upon duplex unwinding in the AGO-small RNA complex (Matranga et al., 2005; Rand et al., 2005). Higher plants encode four distinct DCLs exemplified by arabidopsis DCL1, DCL2, DCL3 and DCL4 (Margis et al., 2006), of which three have clearly defined functions. DCL1 makes miRNAs, mostly 21 nt in size (Park et al., 2002) while DCL3 makes 24-nt siRNAs involved in transcriptional silencing of repetitive elements through DNA methylation (Xie et al., 2004). DCL4 makes different types of 21-nt siRNAs, most importantly the bulk of antiviral siRNAs and endogenous siRNAs from dsRNA synthesized by the RNA-dependent RNA Polymerase RDR6 following initial targeting of a precursor transcript by one or more miRNAs (Allen et al., 2005; Dunoyer et al., 2005; Gasciolli et al., 2005; Xie et al., 2005; Yoshikawa et al., 2005; Bouche et al., 2006; Deleris et al., 2006). The exact functions of DCL2 remain less clearly defined. Biochemically, DCL2 is distinct from other DCLs in two regards. First, it produces 22-nt siRNAs, different from the 21-nt and 24-nt size classes produced by other plant DCLs (Xie et al., 2004). This could have biological significance, because the 22-nt size class has a unique, and so far not clearly explained, ability to trigger dsRNA production and siRNA amplification via RDR6 (Chen et al., 2010; Cuperus et al., 2010). Indeed, DCL2 is required for RDR6-dependent siRNA amplification in transgene silencing, and inactivation of *DCL4* greatly stimulates this process, probably because DCL2 has no competition for transgene dsRNA in the absence of DCL4 (Mlotshwa et al., 2008; Parent et al., 2015). Second, in contrast to other DCLs, its biochemical activity has only been observed indirectly through absence of 22-nt siRNAs in *dcl2* knockout mutants (Xie et al., 2004; Qi et al., 2005; Blevins et al., 2006; Bouche et al., 2006; Deleris et al., 2006; Nagano et al., 2014), perhaps suggesting lower enzymatic activity than that of other DCLs, or that it requires labile co-factors that preclude detection of activity in cell-free systems. Nonetheless, a large body of evidence indicates that DCL2 has important biological functions that manifest themselves in several ways, so far regarded as distinct. First, together with DCL4, it is required for basal antiviral resistance, i.e. for the defense against viruses not adapted to specific hosts by employment of virulence factors capable of suppressing the host antiviral RNAi response (Deleris et al., 2006; Wang et al., 2011; Andika et al., 2015). Experimentally, such a setting is typically mimicked by inactivation of anti-RNAi virulence factors of an adapted virus. The requirement for DCL2 in basal antiviral resistance is observed both at local infection sites, and systemically (Deleris et al., 2006). Second, DCL2 is particularly important for systemic RNAi (Taochy et al., 2017; Chen et al., 2018), an activity that may be linked to its alleviation of symptoms even in infections with adapted viruses (Zhang et al., 2012). Third, in the absence of DCL4 or certain RNA decay factors, DCL2 confers a developmental phenotype with delayed post-embryonic growth and leaf anthocyanin accumulation and/or yellowing (Parent et al., 2015; Zhang et al., 2015). This phenotype is strongly exacerbated by simultaneous mutation of *DCL4* and *DCL1* (Bouche et al., 2006), and in certain combinations of *dcl4* and mutants in RNA decay, it becomes outright lethal at the embryonic stage (Zhang et al., 2015).

At present, explanations for all of the important biological roles of DCL2 rely on the assumption that their biochemical basis is the production of siRNAs that silence complementary targets through the action of RISC. In basal antiviral resistance, DCL2-mediated 22-nt siRNAs can indeed be detected, albeit only when DCL4 is inactivated, such that disappearance of both 21-22-nt siRNAs in *dcl2 dcl4* mutants correlates with complete loss of basal antiviral resistance (Deleris et al., 2006; Wang et al., 2011). Nonetheless, even if the 22-nt siRNAs produced by DCL2 could in theory account for DCL2-mediated antiviral resistance, previous observations of partial loss of basal antiviral resistance in the presence of DCL2-dependent 22-nt siRNAs upon inactivation only of DCL4 may also be interpreted to mean that the siRNAs produced by DCL2 are insufficient to confer resistance (Deleris et al., 2006; Wang et al., 2011; Andika et al., 2015). In systemic RNAi, the 22-nt siRNAs produced by DCL2 are proposed to condition efficient amplification of siRNAs via RDR6 in both source and recipient tissues, probably via properties of RISC containing 22-nt siRNA (Taochy et al., 2017). Finally, the severe growth phenotypes conferred by DCL2 in the absence of DCL4 and/or in RNA decay mutants are proposed to result from ectopic silencing by DCL2-dependent 22-nt siRNAs, potentially because of their ability to engage the small RNA amplification machinery in an uncontrolled way. Consistent with this hypothesis, ectopic silencing by 22-nt siRNAs of the nitrate reductase-encoding *NIA1/2* and of *SMXL4/5* encoding phloem differentiation factors is observed in *dcl4* mutants, and knockout of these genes recapitulates some of the DCL2-dependent defects observed in *dcl4* mutants (Zhang et al., 2015; Wu et al., 2017). Clear proof that silencing of *SMXL4/5* and/or *NIA1/2* by the ectopic 22-nt siRNA populations actually causes the growth defects in *dcl4* mutants has not been reported, however, as this would require phenotypic analysis upon specific reversion of silencing of these genes in *dcl4* mutant backgrounds; a condition that is difficult to achieve experimentally. Thus, at present it cannot be regarded as established fact that *SMXL4/5* or *NIA1/2* silencing underlies the DCL2-dependent growth defects observed in *dcl4* mutants. Indeed, the fact that the presence of DCL2 in some genetic backgrounds (e.g. *dcl4 ski2*, *dcl4 sgt1b*, or *xrn4 ski2* (Zhang et al., 2015; Nielsen et al., 2023) can give rise to phenotypes much stronger than the defects observed, for instance, in *smxl4/5* knockout mutants (Zhang et al., 2015; Wu et al., 2017) suggests, as a minimum, that additional relevant targets of ectopic DCL2-dependent siRNAs remain to be identified. Alternatively, and similar to the situation described for the role of DCL2 in basal antiviral resistance, the DCL2-dependent siRNA populations may not, as a matter of fact, cause the severe DCL2-dependent developmental phenotypes. In principle, answers to this fundamental question should be tangible by rigorous mutational analysis of the *DCL2* gene: if *DCL2* has functions distinct from mere production of siRNAs guiding RISC to silence complementary targets, it should be possible to isolate separation-of-function alleles that either maintain siRNA production, yet lose other DCL2 functions, or lose siRNA production while maintaining other DCL2 functions. Here, we report the engineering of a series of DCL2 point mutants affecting the catalytic activity of RNAseIII domains and the ATP-binding/hydrolysis activity of the helicase domain, as well as 10 novel alleles of *DCL2* isolated by a forward genetic screen for suppressors of DCL2-dependent growth inhibition. The results show that while all reported functions of DCL2 rely on RNaseIII catalytic activity, separation-of-function alleles that allow wild type levels of 22-nt siRNAs to accumulate can be identified. Surprisingly, simple reduction of DCL2 dosage in plants heterozygous for a *DCL2* wild type allele is also sufficient to achieve similar separation of function. These results underscore the need to revise current models for DCL2 function, and we discuss them in the light of findings in the accompanying paper that reveal a function of DCL2 in activation of innate immunity in response to cytoplasmic double-stranded RNA (Nielsen et al., 2023).

## RESULTS

### Growth defects occur upon loss of DCL4 protein and depend on DCL2 dosage and dsRNA

A key motivation to undertake the present work stems from our observation that when young wild type and *dcl4* mutant arabidopsis seedlings were transferred from agar plates to soil, but not when germinated directly in soil, *dcl4* knockout mutants often exhibited growth defects and chlorosis (**Figure 1A,B**), albeit with incomplete penetrance and batch-to-batch variability in the exact frequency of phenotypic penetrance. Similar growth defects in *dcl4* mutants have been noted by others (Gasciolli et al., 2005; Parent et al., 2015; Wu et al., 2017). To analyze whether the incompletely penetrant growth phenotype of *dcl4* knockout mutants was caused by loss of a specific class of DCL4-dependent siRNA, we first used an allelic series of *dcl4* mutants in two different genetic backgrounds (Col-0 and L*er*) including insertion, non-sense and missense alleles that specifically abolish the production of only certain types of DCL4-dependent siRNAs (**Figure 1C**) (Dunoyer et al., 2005; Xie et al., 2005; Dunoyer et al., 2007; Liu et al., 2012b; Montavon et al., 2018). These analyses established that the growth phenotype was only observed at an appreciable frequency upon loss of DCL4 protein (i.e. in insertion and non-sense mutants; **Figure 1A-C**). We used an artificial miRNA targeting DCL2 (amiR-DCL2) to verify that the growth defects observed in mutants homozygous for the non-sense allele *dcl4-15* in accession L*er* were DCL2-dependent (**Figure 1B**). This was confirmed with the knockout allele *dcl2-1* (Xie et al., 2004) introduced into *dcl4-2t*, both in accession Col-0 (**Figure 1A**). Notably, the G587D substitution in *dcl4-8* that causes loss of all classes of DCL4-dependent siRNAs and gives rise to an enzyme that remains bound to dsRNA substrates (Montavon et al., 2018) did not produce growth inhibition and chlorosis phenotypes (**Figure 1A**), suggesting that access of DCL2 to dsRNA is required to trigger these phenotypes. In addition, we observed that the penetrance of leaf yellowing and growth inhibition was 100% in the *dcl4-2t dcl1-11* double mutant (accession Col-0), and noticed that a *dcl2-1* knockout allele even in the heterozygous state was sufficient to cause substantial suppression of this phenotype (**Figure 1D**). Finally, we verified previous results (Zhang et al., 2015) that inactivation of *RDR6* fully suppressed the growth inhibition and chlorosis phenotypes in *dcl4* knockout mutants (**Figure 1E**). These initial observations suggest that loss of DCL4 protein, not only activity, causes growth defects that depend on dsRNA availability and accessibility, and DCL2 dosage.

**Figure 1.**
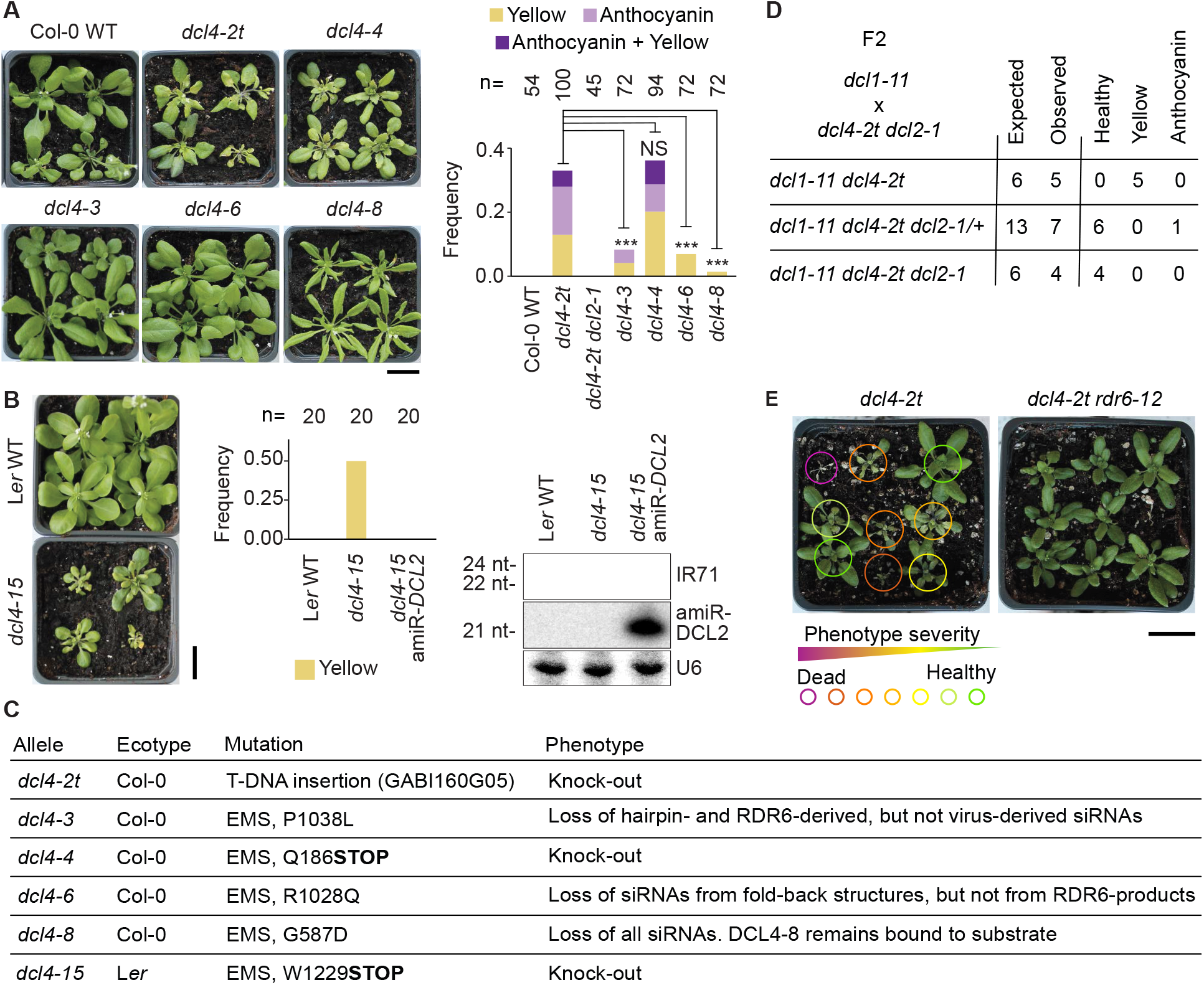
Growth phenotypes of *dcl4* mutants require loss of DCL4 protein, full DCL2 dosage and RDR6. **A**, Rosette phenotypes of 4-week old plants of the indicated genotypes. Plants were grown in long days, first on sterile MS plates, and healthy seedlings at day 11 were transferred to soil and grown for 17 additional days. Left, representative images. Right, fraction of plants showing either leaf yellowing (yellow), visible anthocyanin accumulation (anthocyanin), or both (yellow + anthocyanin). Asterisks indicate significance of difference compared to *dcl4-2t* (not calculated for the obvious cases of Col-0 WT and *dcl4-2t dcl2-1*), ***, P < 0.001 (χ^2^ test); NS, not significant. **B**, Right and middle panels, rosette phenotypes of the indicated genotypes. Growth conditions, photographs and quantifications as described in A. Right panel, small RNA blot shows the expression of the artificial miRNA targeting DCL2 (amiR-DCL2), and the disappearance of the DCL2-dependent 22nt-siRNAs derived from IR71 in the *dcl4-15 amiR*-*DCL2* line. **C**, List of *dcl4* alleles used in this study. **D**, Results of a complete genotyping (for *dcl1-11*, *dcl4-2t* and *dcl2-1* alleles) and phenotyping of an F2 population of 414 individuals resulting from a cross of *dcl1-11* to *dcl4-2t dcl2-1*. The observed individuals of each genotype are sorted according to the phenotypic categories described in A. **E**, Representative rosettes of *dcl4-2t* and *dcl4-2t rdr6-12* plants grown as described in A. For *dcl4-2t*, plants are circled in colors that indicate the severity of growth arrest. Size bars, 2 cm.

### RNaseIII and helicase activities of DCL2 are required for growth and antiviral defense phenotypes

As a first step to define the biochemical basis of the function of DCL2 in causing growth phenotypes and antiviral defense, we engineered point mutants in the RNaseIII catalytic site (DCL2^E1102A^) or in the DExH ATP-binding site (DCL2^D152N^ and DCL2^E153Q^). We then tested stable, transgenic lines expressing these DCL2 mutants in the *dcl4-2t dcl2-1* double knockout background for phenotypic effects and ability to produce 22-nt siRNAs. None of those lines exhibited growth phenotypes (**Figure 2A**), and DCL2-dependent siRNA generation was either completely abolished (catalytic RNaseIII mutant) or severely reduced (**Figure 2B**). Similarly, resistance to turnip crinkle virus deprived of its silencing suppressor, P38 (TCVΔP38, (Qu et al., 2008)), to mimic infection by a non-adapted virus, was compromised in transgenic lines expressing these DCL2 mutant proteins in *dcl4 dcl2* knockout backgrounds (**Figure 2C**). Thus, the dicer activity of DCL2 is required both to confer growth phenotypes in the absence of DCL4, and to confer basal antiviral resistance.

**Figure 2.**
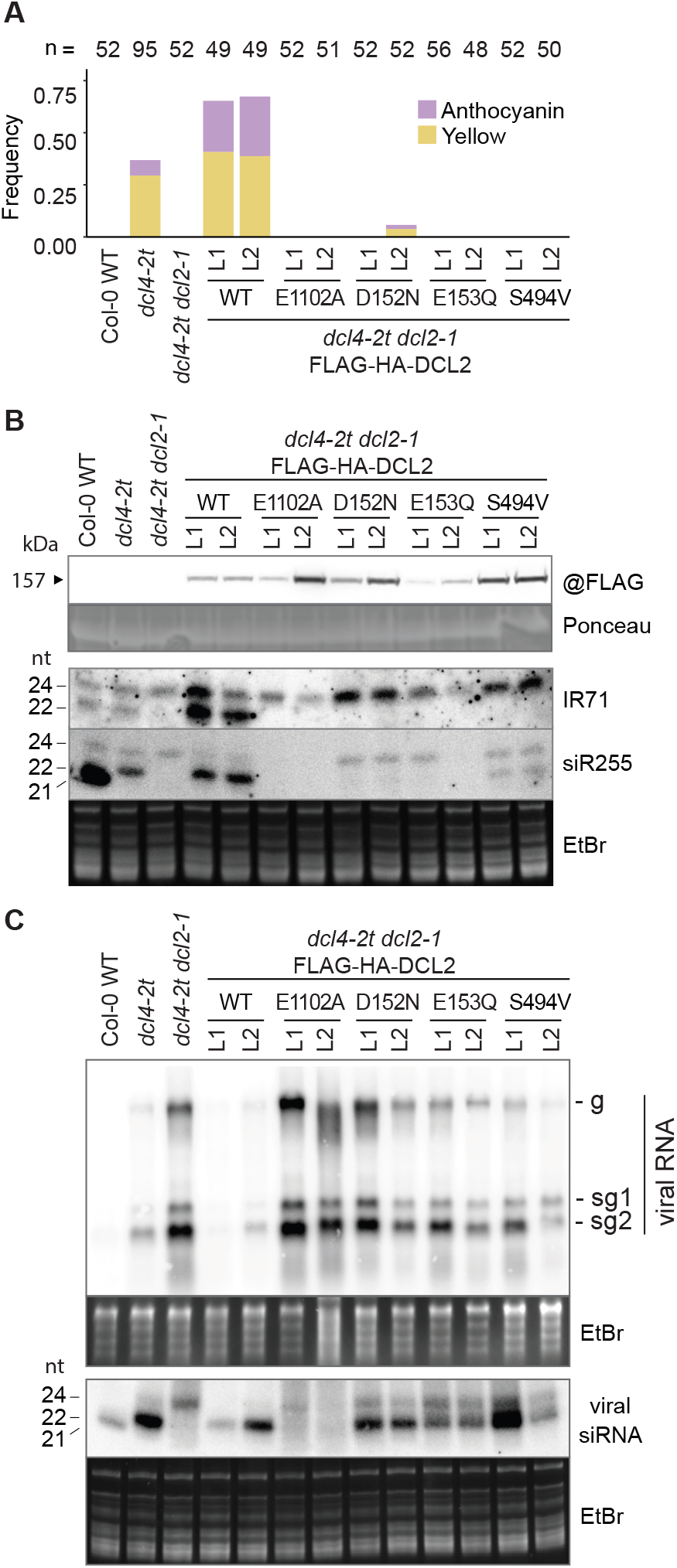
RNaseIII and helicase activities are necessary for Dicer activities and biological functions of DCL2. **A**, Quantification of rosette phenotypes of the indicated genotypes as in Figure 2A. L1 and L2 denote independent transgenic lines. **B**, Molecular analyses of 14-day old sterile-grown seedlings with no apparent phenotypes of the lines used in A. Top, protein blot of total lysates developed with anti-FLAG antibodies. Ponceau staining of the membrane is shown as loading control. Bottom, small RNA blots hybridized to radiolabeled probes complementary to endogenous siRNAs. IR71, inverted repeat 71; siR255, a tasiRNA. Hybridizations were carried out consecutively to the same membrane after probe stripping. EtBr staining of the upper part of the gel is shown as loading control. **C**, RNA blots showing accumulation of viral RNA species upon infection of plants of the indicated genotypes with TCVΔP38. Inoculated leaves were analyzed at 5 days post-inoculation. Top, RNA blot of total RNA hybridized to a radiolabeled probe complementary to TCV. g, genomic RNA; sg, subgenomic RNAs. EtBr staining of the gel is shown as loading control. Bottom, small RNA blot hybridized to a radiolabeled probe complementary to TCV. EtBr staining of the upper part of the gel is shown as loading control.

We also included the helicase mutant S494V in this initial mutational analysis, because the analogous A-V substitution in *Drosophila* Dcr-2 causes defective induction of a defense gene in response to viral infection despite abundant virus-derived siRNA accumulation (Deddouche et al., 2008). Thus, the S494V mutant may be regarded as a candidate separation-of-function mutant. This mutant behaved largely similarly to the ATP-binding site mutants of the helicase domain with suppression of growth phenotypes, defective basal resistance to TCVΔP38, and severely reduced endogenous 22-nt siRNA production, in particular from the endogenous hairpin IR71 (**Figure 2A-C**). However, in contrast to the catalytic RNaseIII and DExH helicase mutants, the DCL2^S494V^ mutant showed detectable accumulation of the RDR6-dependent siR255, and in at least one of the lines examined, TCV-derived siRNAs were abundant despite reduced antiviral resistance (**Figure 2C**).

A number of conclusions and inferences follow from these initial mutational analyses. First, the results obtained with the lines expressing the catalytically inactive DCL2^E1102A^ mutant show that catalytic activity is required for both DCL2 functions examined. We also infer that dsRNA binding by DCL2 is not sufficient to cause growth inhibition and antiviral resistance, since the domains known to be required for dsRNA binding in DCL1 (Wei et al., 2021) remain intact in the catalytic mutants, and since the disappearance of 24-nt viral siRNA in DCL2^E1102A^ (**Figure 2C**) suggests dominant negative influence on other DCLs, consistent with dsRNA binding. Second, we conclude that the helicase activity of DCL2 is required for siRNA production, perhaps suggesting that DCL2 acts processively and that the ATP hydrolysis cycle of the helicase domain is required for translocating the dsRNA between two catalysis events, as reported for DCL1 (Liu et al., 2012a; Wei et al., 2021). Third, although the residual siRNA-producing activity detected in the DCL2^S494V^ mutant cannot be regarded as a strong argument for separation of function in and of itself, it clearly motivates further efforts to investigate the feasibility of separation of function, because of proficiency plants expressing DCL2^S494V^ for some level of virus-derived siRNA production, yet inability to confer full anti-viral resistance.

### Contrasting effects of DCL2 dosage on siRNA accumulation and growth inhibition

We next returned to the observation that growth phenotypes of *dcl4 dcl1* could be suppressed by *dcl2-1* in the heterozygous state. Indeed, *dcl4-2t dcl2-1/+* also showed significantly reduced penetrance of growth phenotypes compared to *dcl4-2t* (**Figure 3A**). Remarkably, heterozygosity of *DCL2* also compromised TCVΔP38 resistance in combination with inactivation of *DCL4* (**Figure 3B**). Because of the incomplete penetrance of *dcl4-2t* single mutants, we sought to corroborate the suppression of growth phenotypes by *DCL2* heterozygosity in a genetic background in which the DCL2-dependent growth phenotype is fully penetrant. For this analysis, we chose the *dcl4 sgt1b* double mutant, shown in the accompanying paper to exhibit strong and fully penetrant DCL2-dependent growth inhibition (Nielsen et al., 2023). Compared to *dcl4-2t dcl1-11* that also exhibits full penetrance of growth inhibition, *dcl4 sgt1b* is easier to handle as it does not suffer from the severely reduced fertility and extremely slow growth of *dcl4-2t dcl1-11*. Both *dcl2-1* and the non-sense *dcl2*-*11* allele (**Table S1**, see below) substantially suppressed *dcl4 sgt1b* in the heterozygous state (**Figure 3C-E**). Further analysis of a series of heterozygous *dcl2* point mutants (**Table S1**, see below) yielded similar results (**Figure S1**). Importantly, this semi-dominance of *dcl2* mutant alleles was not paralleled by a defect in siRNA production. *dcl4 dcl2-1/+* seedlings accumulated levels of endogenous hairpin siRNAs (IR71) similar to *dcl4* (**Figure 4A**), and small RNA-seq of sterile-grown, asymptomatic seedlings did not reveal reduced 22-nt siRNA levels transcriptome-wide in *dcl4-2t* mutants heterozygous for *dcl2-1* compared to *dcl4-2t* mutants homozygous for the *DCL2* wild type allele (**Figure 4B**). This was also true for several individual genes whose ectopic silencing in *dcl4* was previously suggested to play a role in inducing DCL2-dependent growth phenotypes (**Figure 4C**). These results show that full DCL2 dosage is required both for basal antiviral resistance and to induce growth phenotypes in the absence of *DCL4*, but not to reach wild type levels of 22-nt siRNAs.

**Figure 3.**
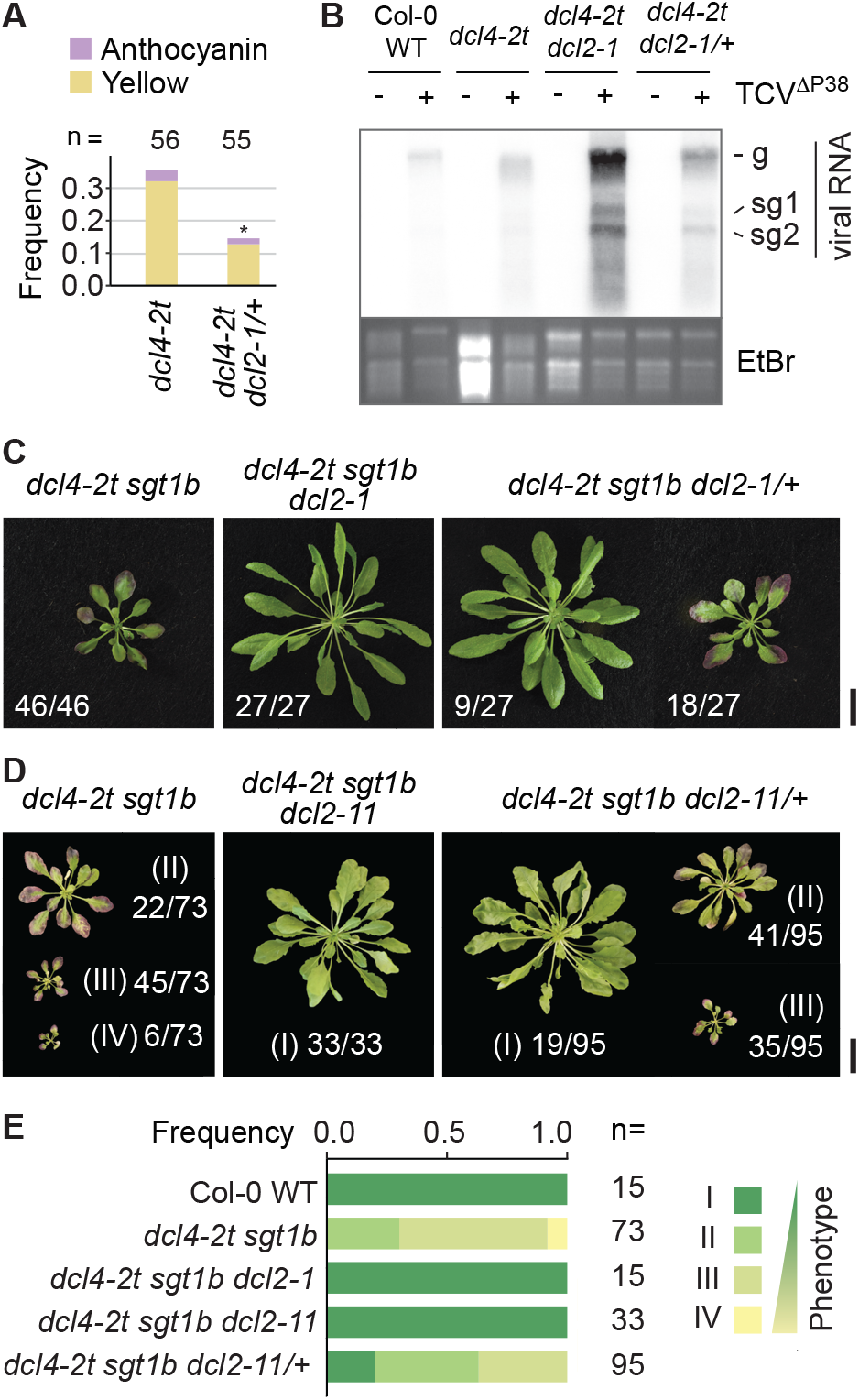
Full DCL2 dosage is required for DCL2 to induce growth defects and for basal antiviral defense. **A**, Quantification of rosette phenotypes of the indicated genotypes as in Figure 2A. *, significance of difference P < 0.05 (χ^2^ test). **B**, RNA blot of total RNA isolated from leaves either mock-treated or inoculated with TCVΔP38 RNA at 5 days post-inoculation. The blot was hybridized to a radiolabeled probe complementary to TCV. g, genomic RNA; sg, subgenomic RNAs. EtBr staining of the gel prior to blotting is shown as loading control. **C**, Rosette phenotypes of the indicated genotypes. Fractions below each photograph indicate the number of individuals in the inspected population exhibiting a phenotype corresponding to the photographed individual, divided by the total number of inspected plants. Size bar, 2 cm. **D**, Phenotypic categorization of rosettes of 5-week old plants grown under short-day conditions on soil. Category I corresponds to green, healthy rosettes, and three categories of different phenotypic severity (II, III, IV) observed in *dcl4-2t/sgt1b* plants are shown. **E**, Representative photographs of the rosette phenotypes quantified in C. Fractions indicate number of plants in each category divided by the total number of plants inspected for each genotype. All genotypes involving dcl2 in the heterozygous state were confirmed by PCR (see Materials and Methods). Scale bars, 1 cm.

**Figure 4.**
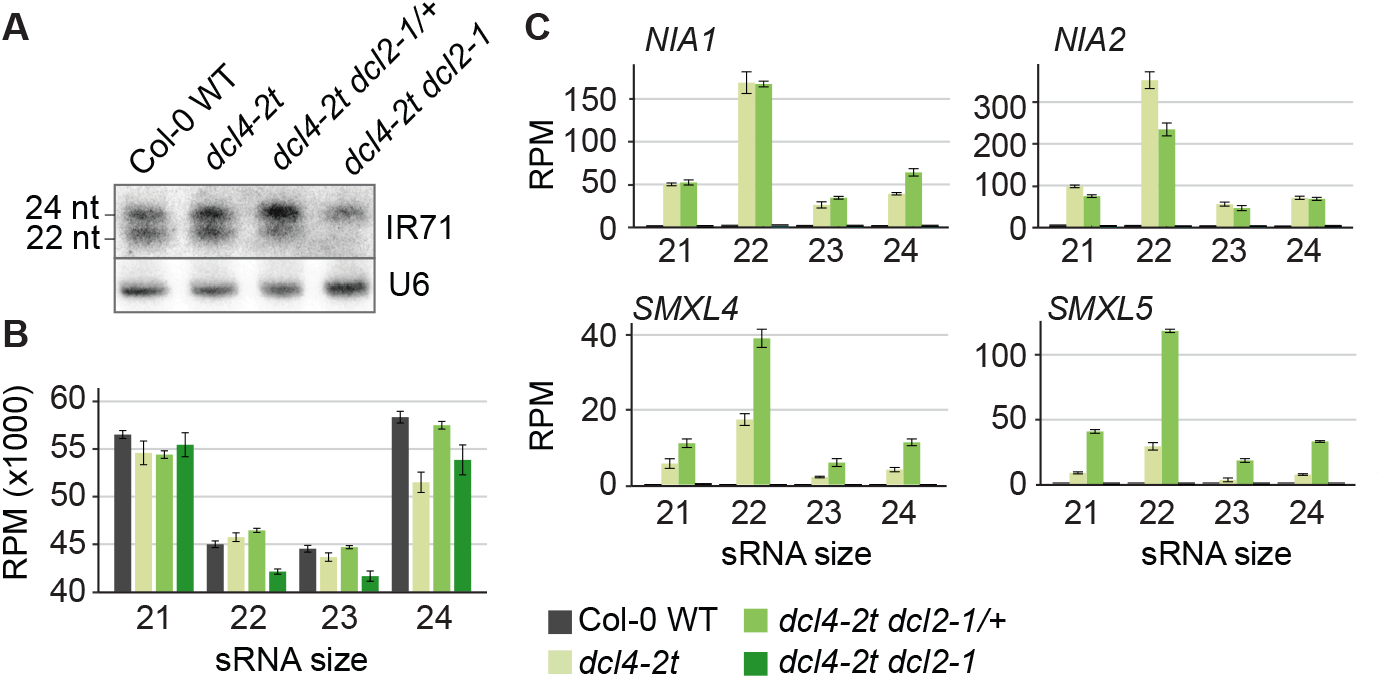
Full DCL2 dosage is not required for siRNA production. **A**, Small RNA blot with seedling RNA hybridized consecutively to radiolabeled probes complementary to IR71 and, as a loading control, to U6 snRNA. **B**, Small RNA-seq conducted on RNA isolated from 21-day old seedlings of the four indicated genotypes. Total reads per million with insert sizes of 21-24 nt are shown. **C**, Normalized number of small RNA reads of the indicates sizes mapping to the genes *NIA1*, *NIA2*, *SMXL4* and *SMXL5*.

### DCL2-dependent growth inhibition is not caused by silencing siRNAs

To corroborate the important conclusion that silencing via ectopic DCL2-dependent siRNAs does not cause growth inhibition, we conducted a forward genetic screen for suppressors of DCL2-dependent growth phenotypes in *dcl4 sgt1b*, in this case with an eye towards recovery of informative mutant alleles of *DCL2*. We identified 10 mutant alleles of *DCL2*, including several non-sense alleles, as expected (**Figure 5A, Table S1**). Four point mutations clustering in the ATP-binding domain of the helicase (*dcl2-6* to *dcl2-9*) and one in the PAZ domain (*dcl2-12*) were investigated further. These mutations suppressed the growth phenotype (**Figure 5B**), but did not abrogate the production of an endogenous 22-nt DCL2-dependent siRNA in the absence of DCL4, as assessed by RNA blot (**Figure 6A**). Since *dcl2-12* did not fully suppress the *dcl4 sgt1b* growth phenotype (**Figure 5A**), it may simply be a weak mutant allele of *DCL2* that reduces its different functions partially, but evenly. In contrast, the mutants located in the helicase conferred strong suppression of *dcl4 sgt1b* growth phenotypes. Because *dcl2-8* was least affected for accumulation of the endogenous siRNA, siR255 (**Figure 6A**), we selected it for small RNA-seq analysis. This analysis showed that in contrast to the *dcl2-1* knockout mutant, the accumulation of 22-nt siRNAs in *dcl2-8* mutants was similar to wild type (**Figure 6B**). In addition, the resistance to TCVΔP38 was compromised in *dcl2-8* and in several other point mutants, despite the fact that infection triggered abundant 22-nt virus-derived siRNA production, most notably in *dcl2-8* and *dcl2-12* (**Figure 6C,D**). We conclude that DCL2-dependent growth defects can be suppressed by point mutations in *DCL2* despite steady-state siRNA profiles similar to those of plants with wild type *DCL2*. Similarly, defective antiviral responses can be observed despite substantial production of DCL2-dependent virus-derived siRNAs. We note, however, that even if near-wild type levels of DCL2-dependent siRNAs accumulate in uninfected *dcl2-8* mutants *in vivo*, the mutant protein is unlikely to exhibit full catalytic activity, as judged by the relative amounts of viral gRNA and 22-nt virus-derived siRNAs in wild type and mutant (**Figure 6C,D**). We could not directly test the biochemical activity of this and other DCL2 mutant proteins, because contrary to DCL4, immunopurified DCL2 did not exhibit detectable activity towards a dsRNA substrate *in vitro* (**Figure S2**). In summary, the results of analyses of plants with reduced DCL2 dosage or with specific point mutations in DCL2 indicate that DCL2-dependent growth restriction, and probably even antiviral resistance at local infection foci, do not rely on silencing by siRNAs. Intriguingly, however, the results also suggest that full catalytic activity of DCL2 is required for both of these DCL2-dependent effects, an intricate point that we treat in some detail in the Discussion.

**Figure 5.**
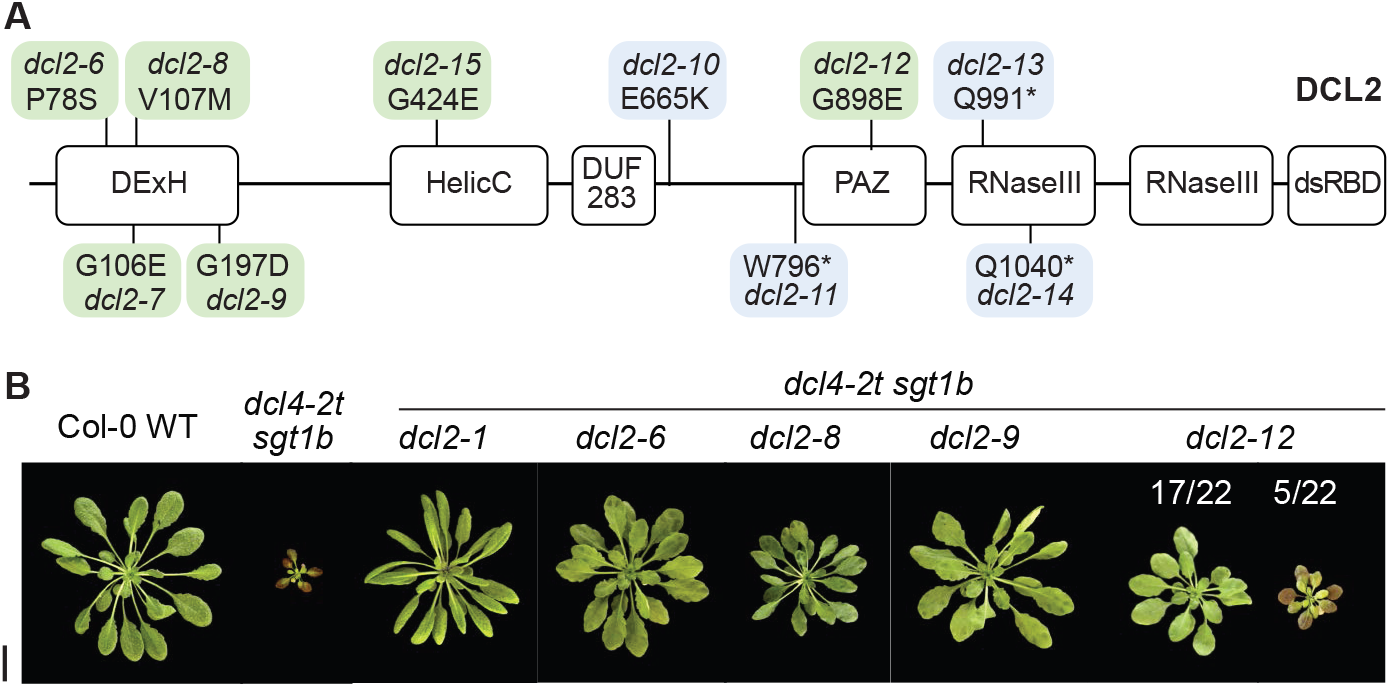
Phenotypes of *dcl2* point mutants that suppress growth defects in *dcl4/sgt1b*. **A**, Schematic overview of *dcl2* mutant alleles recovered by forward genetic screening for *dcl4-2t/sgt1b* suppressors. Green, missense mutations resulting in at least partial Dicer activity; blue, missense or non-sense mutations resulting in complete loss of Dicer activity. Domain designations: ATP, DExH and HelicC, helicase domains; DUF283, Dicer dsRNA-binding fold; PAZ, Piwi-Argonaute-Zwille domain; RNase III, RNAse III domains; dsRBD, dsRNA-binding domain. See also Table S1. **B**, 49-day-old rosettes of the indicated genotypes grown under short day conditions. Scale bar, 2 cm. See also Figure S1.

**Figure 6.**
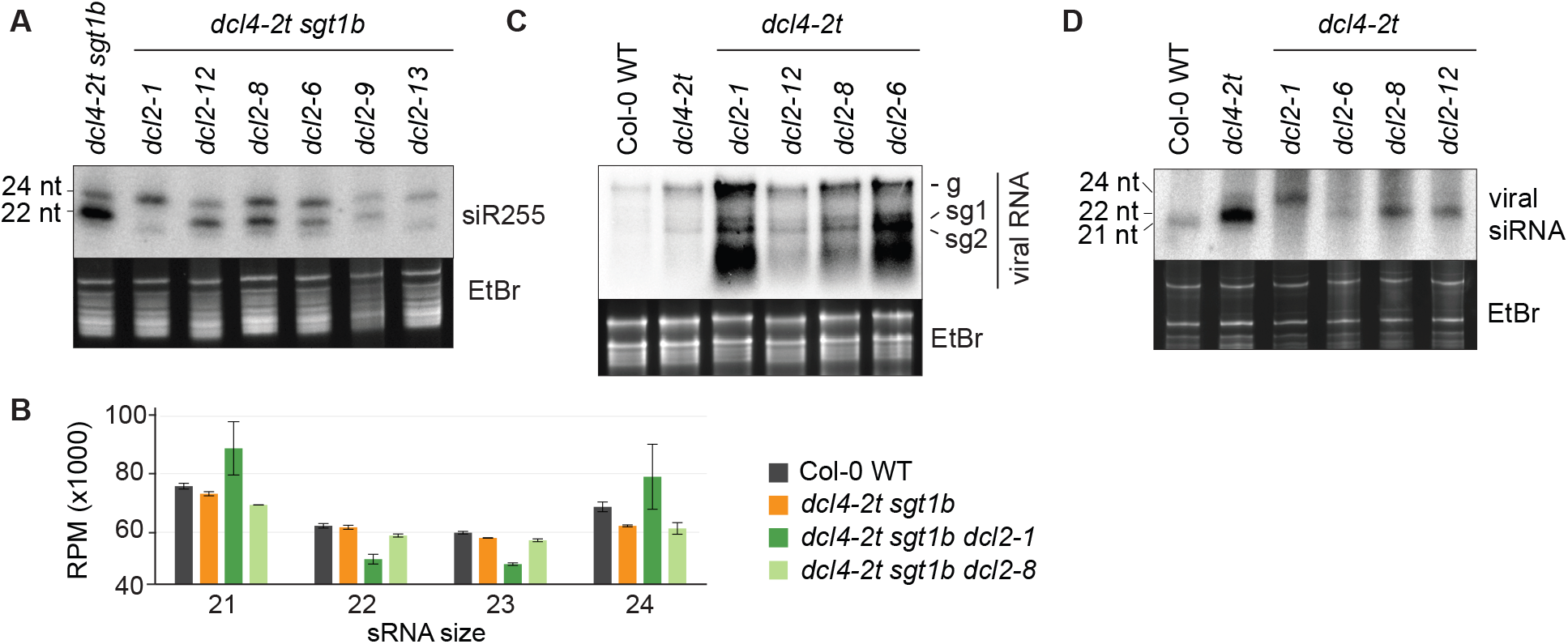
Small RNA accumulation and defective antiviral resistance in *dcl2* point mutants. **A**, Small RNA blot hybridized to a radiolabeled probe complementary to the tasiRNA siR255. EtBr staining of the upper part of the gel is shown as loading control. **B**, Normalized total small RNA counts, sorted by small RNA size, of the indicated genotypes. Small RNA-seq was performed from total RNA extracted from 21-day-old seedlings. **C**, Accumulation of viral gRNA in TCVΔP38-infected leaves at 5 days post-inoculation, assessed by RNA blot. **D**, Small RNA blot showing TCV-derived siRNAs in the same leaves. g, genomic RNA; sg, subgenomic RNAs. EtBr staining is shown as loading control. See also Figure S2.

## DISCUSSION

### Incoherence between biology and known biochemistry of DCL2: implications and solutions

The results of this study strongly indicate that two major effects of DCL2, growth restriction in the absence of DCL4 and basal antiviral resistance together with DCL4 in inoculated tissues, do not depend on silencing by DCL2-dependent siRNAs. The argument relies on the fact that both effects can be suppressed in genetic backgrounds, either homozygous point mutants of *DCL2* or knockout alleles of *DCL2* in the heterozygous state, that produce substantial amounts of DCL2-dependent siRNAs. Indeed, the fact that the siRNA profiles of *dcl4-2t dcl2-8* and *dcl4-2t dcl2-1/+* are similar to that of *dcl4-2t*, yet phenotypic suppression is substantial (for *dcl4-2t dcl2-1/+*) or complete (for *dcl4-2t dcl2-8*), all but excludes the possibility that DCL2-dependent siRNAs underlies growth phenotypes upon loss of DCL4. This conclusion is all the more surprising, because thus far, biological effects of plant DCL enzymes have been explained by their requirement for RNAi through siRNA production, with the only important exception being participation of DCL4 in co-transcriptional transcript cleavage to bring about transcription termination at the *FCA* locus (Liu et al., 2012b). The results have a number of implications, and beg at least three questions that we discuss below.

First, are any previously reported experimental observations consistent with the conclusions reached here? Indeed. In one case, Wang and co-authors analyzed the basal antiviral resistance to a cucumber mosaic virus variant carrying an inactivating mutation in the 2b reading frame encoding its RNAi suppressor. *dcl2* single mutants did not differ in susceptibility from wild type, but in *dcl4* single mutants, partial loss of resistance despite abundant DCL2-dependent 22-nt siRNAs was observed. This led the authors to conclude that “*The 21-Nucleotide, but Not 22-Nucleotide, Viral Secondary Small Interfering RNAs Direct Potent Antiviral Defense by Two Cooperative Argonautes in Arabidopsis thaliana”* (Wang et al., 2011), in direct support of the conclusions reached here. Similarly, inactivation of *DCL4*, but not of *DCL2*, is sufficient to break basal resistance in arabidopsis leaves inoculated with potato virus X (Andika et al., 2015). One may also argue, however, that the additional loss of resistance observed upon mutation of *DCL2* correlates with disappearance of the 22-nt siRNAs, and thereby constitutes an argument in favor of their antiviral activity. Such a counterargument would have little weight in our case, where the 22-nt siRNA populations are similar in genetic backgrounds with contrasting phenotypes.

Second, do our results imply that 22-nt silencing RNAs produced by DCL2 are unable to act as guides for RNAi? Not in the least. The evidence for a specific relevance of DCL2-dependent 22-nt siRNAs in amplified transgene silencing is clear (Mlotshwa et al., 2008; Parent et al., 2015), and is almost certainly a manifestation of activity of RISC loaded with the 22-nt siRNAs (Sakurai et al., 2021; Vigh et al., 2021; Yoshikawa et al., 2021). Indeed, our results remain compatible with a role of DCL2-dependent siRNAs in some aspects of antiviral defense. The infection assays reported here all focused on the analysis of inoculated leaves, hence local infection sites rather than tissues reached upon systemic movement of the virus. There is good evidence that 22-nt siRNAs produced by DCL2 potentiate systemic RNA silencing (Taochy et al., 2017; Chen et al., 2018), and play roles in defense against adapted viruses (Zhang et al., 2012). We therefore suggest that a major antiviral function of DCL2 at the initial stage of infection involves a function that does not depend on the silencing activity 22-nt siRNAs, while 22-nt virus-derived siRNAs play a major role in immunizing uninfected cells systemically. Furthermore, silencing by endogenous DCL2-dependent siRNAs, such as those causing color variation in the seed coat of soybean (Jia et al., 2020), can clearly take place. Thus, the implications of the results presented here should not be generalized further than what they aim to address: whether silencing by DCL2-dependent siRNAs causes basal antiviral resistance at local infection sites and growth phenotypes upon loss of DCL4.

Third, if silencing by DCL2-dependent siRNAs is not the cause of growth inhibition and basal antiviral resistance, then what is? In the accompanying paper, a simple explanation is provided that re-establishes coherence between the biology and molecular function of DCL2 (Nielsen et al., 2023). Together with specific intracellular immune receptors of the nucleotide-binding-leucine rich repeat type (NLRs), dicing by DCL2 causes activation of innate immune responses, such that DCL2 in wild type plants mediates cytoplasmic sensing of excess dsRNA, for example during a viral infection in which RNAi is incapacitated. This activity “misfires” upon loss of DCL4 such that processing of endogenous dsRNA by DCL2 causes autoimmunity via NLRs that manifests itself as the strong DCL2-dependent growth inhibition phenotypes analyzed here and by others. The genetics of this model is solid: plants with a DCL2-dependent growth phenotype indeed exhibit a classical autoimmune gene expression profile, and growth phenotypes and immunity-related gene expression can be strongly suppressed by inactivation of two *NLR*s. Similar to mutation of *DCL2*, inactivation of those *NLR*s causes loss of basal antiviral resistance when combined with *dcl4*, but not alone (Nielsen et al., 2023). Thus, together, the two studies provide a satisfactory framework to understand the important biological effects of DCL2 without the implication of a direct silencing activity of its siRNA products. Many important mechanistic questions now await answers, one of which is particularly nagging and is directly related to the mutational analysis of *DCL2* presented here: why is the catalytic activity of DCL2 required for all of its functions when the siRNAs it produces are not?

### A conundrum and its possible solution: DCL2 activity, but not siRNAs, is required for activation of immunity

Since this study establishes that DCL2-dependent growth phenotypes are dependent on (i) endogenous dsRNA (produced by RDR6), (ii) accessibility of DCL2 to dsRNA, (iii) the catalytic activity of DCL2, and the accompanying study shows that the growth phenotypes are in fact manifestations of NLR-mediated autoimmunity (Nielsen et al., 2023), the key question becomes the following: how does DCL2 facilitate sensing of dsRNA to induce NLR-mediated immune responses? In the closing paragraph, we offer some thoughts, naturally somewhat speculative at this point, on this important problem.

We exclude the possibility that mere dsRNA binding to DCL2 is sufficient to trigger immune signaling, because the catalytically dead RNaseIII mutant of DCL2 was completely defective in inducing growth phenotypes in *dcl4 dcl2* mutants. Thus, in this regard, dsRNA sensing via DCL2 differs from mechanisms of cytoplasmic dsRNA sensing in mammals mediated by the Dicer-related helicases RIG-I and MDA5 (Rehwinkel and Gack, 2020). This biochemical difference in dsRNA sensing may be related to the higher production of endogenous dsRNA in plants than in animals, by virtue of the existence of RNA-dependent RNA polymerases with crucial functions in generation of endogenous siRNAs (Matthew et al., 2011). Ideally, therefore, to avoid autoimmunity, plants would require a sensor system that switches on immune responses only when dsRNA is present in such quantities that DCL2 produces a high flux of siRNAs. Such a kinetic sensor system may explain why reduced DCL2 dosage and mutants with reduced activity that nonetheless produce steady state siRNA levels similar to wild type exhibit defects in immune activation. It is at present unclear whether DCL2 itself is the sensor, a scenario that would require physical association of R proteins with DCL2, or whether siRNA products of DCL2, when made at sufficiently high rates, have properties that allow their perception by other sensors. Thus, sensing at the level of RISC cannot be excluded, a scenario which would conceptually parallel the recently described prokaryotic plasmid-sensing systems that rely on oligomerization of heterodimers of a variant AGO protein and a Toll/Interleukin-1 receptor domain protein upon pervasive base pairing of plasmid-derived, AGO-bound guide RNAs to high-copy plasmids (Koopal et al., 2022).

## ACKNOWLEDGMENTS

Theo Bölsterli, Rene Hvidberg and their teams are thanked for plant care. Lena Bjørn Johansson is thanked for help with seed collection, plating and genotyping in connection with transgenic line and mutant constructions. The Nottingham Arabidopsis Stock Centre (NASC) is thanked for providing arabidopsis T-DNA insertion lines. Patrice Dunoyer (*dcl4-2*, *dcl4-3*, *dcl4-4*, *dcl4-6*, *dcl4-8*) and Caroline Dean (*dcl4-15*) are thanked for providing seeds of arabidopsis *dcl4* mutants. Maria Louisa Vigh is thanked for help with assays of DCL4 and DCL2 activity. Detlef Weigel, Christophe Ritzenthaler, Simon Bressendorff, Diego López-Márquez, Emilie Oksbjerg and Dexter Adams are thanked for critical reading of the manuscript. This work was supported by grants to P.B. from the Independent Research Fund Denmark (Sapere Aude Grant 12-133793), Villum Fonden (Project 13397), and the European Research Council (PATHORISC, ERC-2016-CoG 726417).

## AUTHOR CONTRIBUTIONS

CPSN constructed most lines expressing point mutants of DCL2 in *dcl2 dcl4*, conducted all steps of the *dcl4 sgt1b* suppressor screen, and acquired and analyzed all high-throughput sequencing data. LA-H acquired the first evidence for defense activation in *dcl4* and *dcl4 dcl1* mutants, carried out phenotypic analysis of *dcl4* mutants, noticed the effect of *dcl2* heterozygosity in *dcl4 dcl1* mutants, constructed *dcl4 rdr6* double mutants, initiated work on DCL2 catalytic and helicase mutants, and participated in discussions with PB leading to formulation of the model for DCL2 function proposed here and in the accompanying paper. LH constructed and characterized *dcl4-15*/amiR-*DCL2* lines. NP constructed the DCL2 wild type plasmid used as a starting point for generation of point mutants, and shared the FLAG-HA-DCL4 transgenic line prior to publication. SUA instructed and supervised genetic mapping based on high-throughput sequencing data. PB conceived the project, acquired funding, designed experiments, supervised work, and wrote the manuscript with input from all authors.

## ACCESSION NUMBERS

sRNA sequencing data sets generated in this study were submitted to the European Nucleotide Archive under accession number PRJEB52819.

## MATERIALS AND METHODS

### Plant material and growth conditions

The *dcl4* mutants used (*dcl4-2t*, *dcl4-15*, and *dcl4-3*, *dcl4-4*, *dcl4-6*, *dcl4-8*) and the transgenic line expressing 2xFLAG2x-HA-DCL4 have been described (Dunoyer et al., 2005; Xie et al., 2005; Liu et al., 2012b; Pumplin et al., 2016; Montavon et al., 2018). The SUC:SUL transgene (Dunoyer et al., 2005) was outcrossed from *dcl4-3*, *dcl4-4*, *dcl4-6*, *dcl4-8* by crosses to Col-0 and selection of the relevant *dcl4* mutants by PCR in the F2 generation. The mutants in *sgt1b* (*edm1*), *rdr6* (*rdr6-12*) used in the study, and the *dcl2-1* mutant have also been described (To▫r et al., 2002; Peragine et al., 2004; Xie et al., 2004). Unless otherwise stated, plants were grown in soil (Plugg/Såjord (seed compost), SW Horto) under a long day cycle (day: 16 hours light, 130 μmol m^− 2^ s^−1^, 21°C; night: 8 hours darkness, 18°C), in Percival growth chambers equipped with Philips Master TL-D 36W/840 and 18W/840 bulbs. *A. thaliana* seeds were surface sterilized before stratification and germination. Surface sterilization was performed by treatment with 70% ethanol for 2 minutes, then in 1.5 % NaOCl, 0.05 % Tween-20 for 10 minutes, followed by two washes in deionized water. After surface sterilization seeds were stratified by incubation in the darkness at 4°C for 3 days. Seeds were then germinated on either soil (Plugg/Såjord (seed compost), SW Horto) or on Murashige-Skoog (MS) agar plates (4.4 g/L MS salt mixture, 10 g/L sucrose, 8 g/L agar, pH 5.7).

### Plant genotyping and phenotyping

All T-DNA lines were genotyped using PCR with 2 different primer sets. One primer set to detect the wild type allele with forward and reverse primers on opposite sides of the T-DNA. The other primer set detects the T-DNA insertion allele, and uses either the forward or reverse primer from the first set, together with a primer inside the left border of the T-DNA. For point mutants, derived Cleaved Amplified Polymorphic Sequence (dCAPS) markers (Neff et al., 1998) were designed to allow genotyping by restriction digest following PCR. All primer sets and, if appropriate, cognate restriction enzymes are listed in Table S2. Quantification of incompletely penetrant phenotypes, including the statistical analyses of the observations, was done exactly as described in the accompanying paper (Nielsen et al., 2023).

### Cloning and construction of transgenic lines

The vector for expression of genomic, double FLAG, double HA-tagged version of DCL2 under the endogenous promoter [Pro(DCL2):2xFLAG-2xHA-DCL2^WT^:ter(35S)] was constructed in the pB7m34GW vector by multisite Gateway recombination, as described by (Karimi et al., 2005). Briefly, the DCL2 promoter was cloned by amplifying the 2.6 kb of DNA sequence immediately upstream of the coding sequence start codon, using primers that added gateway recombination sites to enable cloning into a pDONR4-1R plasmid. DCL2 genomic coding sequence was amplified using primers that add gateway recombination sites for cloning into a pDONR2R-3 plasmid. The 2xFLAG-2xHA in pDONR1-2 was a gift of Clement Chevalier. To construct the DCL2 point mutant lines, we used this vector as template for site-directed mutagenesis. Primers were designed according to the Quickchange II manual (Agilent) so that the forward and reverse primers were complementary to each other, and covered the desired mutation site. For D152N we used primers AN5/AN6, for E153Q we used BP11/BP12, for S494V we used CSN116/CSN117, and for E1102A we used BP7/BP8.

The artificial miRNA targeting DCL2 (amiR-DCL2) was designed using the WMD3 – Web MicroRNA Designer (Schwab et al., 2006). A DNA fragment comprising amiR-DCL2 inside the pri-miR319a backbone and flanked by attL1 and attL2 sites (Supplementary Table S3) was synthesized by Integrated DNA Technologies. Using the LR reaction, the pri-amiR-DCL2 was then introduced into a derivative of pGWB502 (Nakagawa et al., 2007) in which the CaMV 35S promoter was replaced by the Cestrum Yellow Leaf Curling Virus CmpC promoter (Stavolone et al., 2003). The resulting construct was then transformed into *dcl4-15* mutants. All primer sequences are listed in Table S2. Plant transformation and selection of transgenic plants and lines were done as described in the accompanying paper (Nielsen et al., 2023).

### Construction and analysis of plants heterozygous for *dcl2* alleles

To construct *dcl4-2t dcl2-1/+*, we emasculated *dcl4-2t dcl2-1* flowers, and used F1 seeds resulting from fertilization with pollen from *dcl4-2t* flowers. After analysis (phenotype counts or TCV ΔP38 infections), the genotypes of individual plants were confirmed to be *dcl4-2t dcl2-1/+* by PCR using the primers listed in Table S2. The *dcl4-2t dcl2-1/+* genotype was also confirmed for plants pooled for RNA extraction for small RNA northern and sequencing analysis. For construction of *dcl4-2t sgt1b* heterozygous for the different *dcl2* alleles (*dcl4-2t sgt1b dcl2-x/+*), we emasculated *dcl4-2t sgt1b dcl2-x*, because these plants were easier to manipulate than *dcl4-2t sgt1b*. Gynoecia of emasculated flowers were pollinated with pollen from *dcl4-2t sgt1b*. F1 plants resulting from these crosses were examined for phenotypes. All F1 plants were genotyped by PCR using the primers listed in Table S2, and confirmed to be *dcl4-2t sgt1b dcl2-x*.

### TCVΔP38 infection and RNA analyses

*In vitro* transcription of TCVΔP38 RNA and infection by hand inoculation of silicon carbide-rubbed leaf surfaces were done exactly as described in the accompanying paper (Nielsen et al., 2023). RNA and protein extractions as well as gel blot analyses of proteins, small RNAs and viral genomic RNAs were also done exactly as described in the accompanying paper (Nielsen et al., 2023).

### Small RNA sequencing and analysis

Libraries for small RNA-seq were prepared from 5 μg of total RNA using the NEBNext Multiplex Small RNA Library Prep Kit for Illumina (E7560S). The quality of libraries and input RNA was assessed using an Agilent 2100 Bioanalyzer. Libraries was sequenced on an Illumina NextSeq 550 platform with a flowcell yielding 75 bp single-end reads. FASTQ files were trimmed using CutAdapt (Martin, 2011), mapped using STAR (Dobin et al., 2013), and numbers of reads mapping to specific genes were counted using featureCounts 1.6.3 (Liao et al., 2014). For the analysis of specific sRNA sizes the sRNA reads were sorted according to size prior to counting, using AWK. Calculations of numbers of reads with specific sizes mapping to all or specific genes, as well as the different plots were constructed in R.

### Dicer assays

A 265 bp *PHABULOSA* fragment in pGEM-T-Easy was PCR amplified with M13 primers to and used for sense and antisene *in vitro* transcriptions with T7 and SP6 polymerases (Promega) in the presence of 1.2 μM of α-^32^P-UTP (10μCi/μL; Hartmann Analytic). Radioactively labeled RNA fragments were PAGE purified, and volumes adjusted so that each contained 1000 cps/μL. Radioactive dsRNA was produced by annealing 1 μL sense to 1 μL antisense RNA in a total volume of 20 μL of 1 x annealing buffer (10 mM Tris-HCl pH 8, 30 mM NaCl, 10 μM EDTA, 1 u Ribolock). The annealing reaction was done in the PCR machine by heating to 75°C for 5 minutes followed by gradual cooling to room temperature in the PCR block for 1 hour without the use of the Pelletier element.

For immuno-purification of 2xFLAG2x-HA-DCL4 (Pumplin et al., 2016) and 2xFLAG-2xHA-DCL2 (this study), we incubated total cleared lysates prepared by mixing 400 mg of inflorescence tissue ground under liquid nitrogen with 1.2 mL of IP buffer with 25 μl of M2 agarose beads (Sigma) for 1 hour at 4°C. Immunocaptured proteins were washed extensively (3-4 times) in IP buffer. During the last wash, immunopurified fractions were split into 2/3 for Dicer assays and 1/3 for FLAG western analysis. For Dicer assays, IP buffer was completely removed with a Hamilton syringe, and 21 μL of 2xTm buffer (133.3 mM KCl, 13.3 mM MgCl_2_, 16.7 mM DTT, 3.3 mM ATP, 0.7 mM GTP, 1% v/v Ribolock) containing 150 cps of annealed dsRNA was added to immunopurified DCL4/DCL2/mock. Reactions were incubated at 25°C for 1 hour, and stopped by addition of 1mL of Trizol, and purified RNA fragments were resolved on a 5% acrylamide:bis 19:1, 7M urea gel in 1xTBE buffer.

### EMS mutagenesis and screen for *dcl4-2t sgt1b* suppressors

Approximately 270 mg *dcl4-2t sgt1b* seeds, an estimated ~14,000 seeds, were incubated with 8 ml of 0.74 % ethyl methanesulfonate (EMS, Sigma) in 0.1 M NaH_2_PO_4_, pH 5.5, 5% dimethyl sulfoxide (DMSO) in 15 ml Falcon tubes in a rotating wheel for 4 hours at room temperature. After 5 washes in 0.1 M Na_2_S_2_O_3_, seeds were dispersed in 0.1% agar and spread directly in soil in families of ~80 individuals. M2 seeds were harvested in pools of ~80 mature M1 plants. About 3000 M2 seeds from each pool were germinated directly in soil and grown under short day conditions (8-hour light, 16-hour dark) in the greenhouse. Plants that did not display the shoot-inhibition phenotype were harvested individually, and the seeds were kept for verification of phenotype and mapping.

### Mapping of EMS mutants

To map the causal mutations of the recovered suppressor mutants, mapping populations were generated by crossing to *dcl4-15 sgt1b* in accession Landsberg. F1 seeds were germinated directly in soil, and F2 seeds obtained by self-pollination were harvested. 400 F2 seeds of each suppressor mutant mapping population were then germinated under short day conditions (8 hour light, 16 hour dark), and flower buds from 30-100 symptomless plants of each F2 population were harvested in nitrogen and pooled. DNA was phenol:chloroform extracted and DNA libraries were prepared using the NebNext Ultra II DNA library kit for Illumina (#E7645S) using the manufacturer’s instructions. The libraries were then sequenced by Novogene Bioinformatics Technology Co., Ltd using paired end 150 bp sequencing with minimum 20 million reads per library. The obtained FASTQ files were trimmed using Cutadapt 2.4 (Adapter sequence: AGATCGGAAGAGC) and read quality was assessed both before and after trimming using FASTQC. We then mapped the reads using Bowtie2 2.2.3 (Langmead and Salzberg, 2012), and used SHOREMAP to obtain the candidate chromosome region and to obtain a list of candidate mutations (Schneeberger et al., 2009).

#### SUPPLEMENTAL MATERIAL

**Supplemental Figure S1**. Suppression of *dcl4-2 sgt1b* phenotypes by DCL2 point mutants allowing siRNA production

**Supplemental Figure S2**. *In vitro* assays of dicer activity of immuno-purified DCL4 and DCL2

**Supplemental Table S1**. Overview of mutant alleles of the arabidopsis *DCL2* gene.

**Supplemental Table S2**. Primers and other oligonucleotides used in the study

## REFERENCES

Allen, E., Xie, Z., Gustafson, A.M., and Carrington, J.C. (2005). microRNA-directed phasing during trans-acting siRNA biogenesis in plants. Cell 121, 207–221.

Andika, I.B., Maruyama, K., Sun, L., Kondo, H., Tamada, T., and Suzuki, N. (2015). Differential contributions of plant Dicer-like proteins to antiviral defences against potato virus X in leaves and roots. The Plant Journal 81, 781–793.

Blevins, T., Rajeswaran, R., Shivaprasad, P.V., Beknazariants, D., Si-Ammour, A., Park, H.S., Vazquez, F., Robertson, D., Meins, F., Jr., Hohn, T., and Pooggin, M.M. (2006). Four plant Dicers mediate viral small RNA biogenesis and DNA virus induced silencing. Nucleic acids research 34, 6233–6246.

Bouche, N., Lauressergues, D., Gasciolli, V., and Vaucheret, H. (2006). An antagonistic function for Arabidopsis DCL2 in development and a new function for DCL4 in generating viral siRNAs. EMBO Journal 25, 3347–3356.

Chen, H.-M., Chen, L.-T., Patel, K., Li, Y.-H., Baulcombe, D.C., and Wu, S.-H. (2010). 22-nucleotide RNAs trigger secondary siRNA biogenesis in plants. Proceedings of the National Academy of Sciences 107, 15269–15274.

Chen, W., Zhang, X., Fan, Y., Li, B., Ryabov, E., Shi, N., Zhao, M., Yu, Z., Qin, C., Zheng, Q., Zhang, P., Wang, H., Jackson, S., Cheng, Q., Liu, Y., Gallusci, P., and Hong, Y. (2018). A Genetic Network for Systemic RNA Silencing in Plants Plant physiology 176, 2700–2719.

Cuperus, J.T., Carbonell, A., Fahlgren, N., Garcia-Ruiz, H., Burke, R.T., Takeda, A., Sullivan, C.M., Gilbert, S.D., Montgomery, T.A., and Carrington, J.C. (2010). Unique functionality of 22-nt miRNAs in triggering RDR6-dependent siRNA biogenesis from target transcripts in Arabidopsis. Nature structural & molecular biology 17, 997–1003.

D’Ario, M., Griffiths-Jones, S., and Kim, M. (2017). Small RNAs: Big Impact on Plant Development. Trends in plant science 22, 1056–1068.

Deddouche, S., Matt, N., Budd, A., Mueller, S., Kemp, C., Galiana-Arnoux, D., Dostert, C., Antoniewski, C., Hoffmann, J.A., and Imler, J.-L. (2008). The DExD/H-box helicase Dicer-2 mediates the induction of antiviral activity in drosophila. Nature Immunology 9, 1425–1432.

Deleris, A., Gallego-Bartolome, J., Bao, J., Kasschau, K.D., Carrington, J.C., and Voinnet, O. (2006). Hierarchical action and inhibition of plant Dicer-like proteins in antiviral defense. Science 313, 68–71.

Ding, S.W., and Voinnet, O. (2007). Antiviral immunity directed by small RNAs. Cell 130, 413–426.

Dobin, A., Davis, C.A., Schlesinger, F., Drenkow, J., Zaleski, C., Jha, S., Batut, P., Chaisson, M., and Gingeras, T.R. (2013). STAR: ultrafast universal RNA-seq aligner. Bioinformatics 29, 15–21.

Dunoyer, P., Himber, C., and Voinnet, O. (2005). DICER-LIKE 4 is required for RNA interference and produces the 21-nucleotide small interfering RNA component of the plant cell-to-cell silencing signal. Nature genetics 37, 1356–1360.

Dunoyer, P., Himber, C., Ruiz-Ferrer, V., Alioua, A., and Voinnet, O. (2007). Intra- and intercellular RNA interference in Arabidopsis thaliana requires components of the microRNA and heterochromatic silencing pathways. Nature genetics 39, 848–856.

Fukudome, A., and Fukuhara, T. (2017). Plant dicer-like proteins: double-stranded RNA-cleaving enzymes for small RNA biogenesis. Journal of Plant Research 130, 33–44.

Gasciolli, V., Mallory, A.C., Bartel, D.P., and Vaucheret, H. (2005). Partially Redundant Functions of Arabidopsis DICER-like Enzymes and a Role for DCL4 in Producing trans-Acting siRNAs. Current biology : CB 16, 1494–1500.

Jia, J., Ji, R., Li, Z., Yu, Y., Nakano, M., Long, Y., Feng, L., Qin, C., Lu, D., Zhan, J., Xia, R., Meyers, B.C., Liu, B., and Zhai, J. (2020). Soybean DICER-LIKE2 Regulates Seed Coat Color via Production of Primary 22-Nucleotide Small Interfering RNAs from Long Inverted Repeats. The Plant cell 32, 3662–3673.

Karimi, M., De Meyer, B., and Hilson, P. (2005). Modular cloning in plant cells. Trends in plant science 10, 103–105.

Koopal, B., Potocnik, A., Mutte, S.K., Aparicio-Maldonado, C., Lindhoud, S., Vervoort, J.J.M., Brouns, S.J.J., and Swarts, D.C. (2022). Short prokaryotic Argonaute systems trigger cell death upon detection of invading DNA. Cell 185, 1471–1486.e1419.

Langmead, B., and Salzberg, S.L. (2012). Fast gapped-read alignment with Bowtie 2. Nat Methods 9, 357–359.

Law, J.A., and Jacobsen, S.E. (2010). Establishing, maintaining and modifying DNA methylation patterns in plants and animals. Nature reviews. Genetics 11, 204–220.

Liao, Y., Smyth, G.K., and Shi, W. (2014). featureCounts: an efficient general purpose program for assigning sequence reads to genomic features. Bioinformatics 30, 923–930.

Liu, C., Axtell, M.J., and Fedoroff, N.V. (2012a). The Helicase and RNaseIIIa Domains of Arabidopsis Dicer-Like1 Modulate Catalytic Parameters during MicroRNA Biogenesis Plant physiology 159, 748–758.

Liu, F., Bakht, S., and Dean, C. (2012b). Cotranscriptional Role for Arabidopsis DICER-LIKE 4 in Transcription Termination. Science 335, 1621–1623.

MacRae, I.J., Zhou, K., and Doudna, J.A. (2007). Structural determinants of RNA recognition and cleavage by Dicer. Nature structural & molecular biology 14, 934–940.

Macrae, I.J., Zhou, K., Li, F., Repic, A., Brooks, A.N., Cande, W.Z., Adams, P.D., and Doudna, J.A. (2006). Structural basis for double-stranded RNA processing by Dicer. Science 311, 195–198.

Margis, R., Fusaro, A.F., Smith, N.A., Curtin, S.J., Watson, J.M., Finnegan, E.J., and Waterhouse, P.M. (2006). The evolution and diversification of Dicers in plants. FEBS Letters 580, 2442–2450.

Martin, M. (2011). Cutadapt removes adapter sequences from high-throughput sequencing reads. EMBnet.journal; Vol 17, No 1: Next Generation Sequencing Data Analysis.

Matranga, C., Tomari, Y., Shin, C., Bartel, D.P., and Zamore, P.D. (2005). Passenger-strand cleavage facilitates assembly of siRNA into Ago2-containing RNAi enzyme complexes. Cell 123, 607–620.

Matthew, R.W., Matthew, W.E., Rebecca, T.C., and Brian, D.G. (2011). The Functions of RNA-Dependent RNA Polymerases in Arabidopsis. The Arabidopsis Book 2011.

Mlotshwa, S., Pruss, G.J., Peragine, A., Endres, M.W., Li, J., Chen, X., Poethig, R.S., Bowman, L.H., and Vance, V. (2008). DICER-LIKE2 Plays a Primary Role in Transitive Silencing of Transgenes in Arabidopsis. PloS one 3, e1755.

Montavon, T., Kwon, Y., Zimmermann, A., Hammann, P., Vincent, T., Cognat, V., Bergdoll, M., Michel, F., and Dunoyer, P. (2018). Characterization of DCL4 missense alleles provides insights into its ability to process distinct classes of dsRNA substrates. The Plant Journal 95, 204–218.

Nagano, H., Fukudome, A., Hiraguri, A., Moriyama, H., and Fukuhara, T. (2014). Distinct substrate specificities of Arabidopsis DCL3 and DCL4. Nucleic acids research 42, 1845–1856.

Nakagawa, T., Suzuki, T., Murata, S., Nakamura, S., Hino, T., Maeo, K., Tabata, R., Kawai, T., Tanaka, K., Niwa, Y., Watanabe, Y., Nakamura, K., Kimura, T., and Ishiguro, S. (2007). Improved Gateway Binary Vectors: High-Performance Vectors for Creation of Fusion Constructs in Transgenic Analysis of Plants. Bioscience, Biotechnology, and Biochemistry 71, 2095–2100.

Neff, M.M., Neff, J.D., Chory, J., and Pepper, A.E. (1998). dCAPS, a simple technique for the genetic analysis of single nucleotide polymorphisms: experimental applications in Arabidopsis thaliana genetics. The Plant Journal 14, 387–392.

Nielsen, C.P.S., Han, L., Arribas-Hernandez, L., Karelina, D., Petersen, M., and Brodersen, P. (2023). Sensing of viral RNA in plants via a DICER-LIKE Ribonuclease. bioRxiv,2023.2001.2010.523395.

Niladri, S., Iwasa, K., Shen, J., Peter, S.,, and Bass, B.L. (2018). Dicer uses distinct modules for recognizing dsRNA termini. Science 359, 329–334.

Parent, J.S., Bouteiller, N., Elmayan, T., and Vaucheret, H. (2015). Respective contributions of Arabidopsis DCL2 and DCL4 to RNA silencing. Plant J 81, 223–232.

Park, W., Li, J., Song, R., Messing, J., and Chen, X. (2002). CARPEL FACTORY, a Dicer homolog, and HEN1, a novel protein, act in microRNA metabolism in Arabidopsis thaliana. Current biology : CB 12, 1484–1495.

Peragine, A., Yoshikawa, M., Wu, G., Albrecht, H.L., and Poethig, R.S. (2004). SGS3 and SGS2/SDE1/RDR6 are required for juvenile development and the production of trans-acting siRNAs in Arabidopsis. Genes & development 18, 2368–2379.

Pumplin, N., Sarazin, A., Jullien, P.E., Bologna, N.G., Oberlin, S., and Voinnet, O. (2016). DNA Methylation Influences the Expression of DICER-LIKE4 Isoforms, Which Encode Proteins of Alternative Localization and Function. The Plant cell 28, 2786–2804.

Qi, Y., Denli, A.M., and Hannon, G.J. (2005). Biochemical specialization within Arabidopsis RNA silencing pathways. Molecular cell 19, 421–428.

Qu, F., Ye, X., and Morris, J.T. (2008). Arabidopsis DRB4, AGO1, AGO7, and RDR6 participate in a DCL4-initiated antiviral RNA silencing pathway negatively regulated by DCL1 Proceedings of the National Academy of Sciences of the United States of America 105, 14732–14737.

Rand, T.A., Petersen, S., Du, F., and Wang, X. (2005). Argonaute2 Cleaves the Anti-Guide Strand of siRNA during RISC Activation. Cell 123, 621–629.

Rehwinkel, J., and Gack, M.U. (2020). RIG-I-like receptors: their regulation and roles in RNA sensing. Nature Reviews Immunology 20, 537–551.

Sakurai, Y., Baeg, K., Lam, A.Y.W., Shoji, K., Tomari, Y., and Iwakawa, H.-o. (2021). Cell-free reconstitution reveals the molecular mechanisms for the initiation of secondary siRNA biogenesis in plants. Proceedings of the National Academy of Sciences 118, e2102889118.

Schneeberger, K., Ossowski, S., Lanz, C., Juul, T., Petersen, A.H., Nielsen, K.L., Jorgensen, J.E., Weigel, D., and Andersen, S.U. (2009). SHOREmap: simultaneous mapping and mutation identification by deep sequencing. Nat Methods 6, 550–551.

Schwab, R., Ossowski, S., Riester, M., Warthmann, N., and Weigel, D. (2006). Highly specific gene silencing by artificial microRNAs in Arabidopsis. The Plant cell 18, 1121–1133.

Song, M.-S., and Rossi, J.J. (2017). Molecular mechanisms of Dicer: endonuclease and enzymatic activity. Biochemical Journal 474, 1603–1618.

Song, X., Li, Y., Cao, X., and Qi, Y. (2019). MicroRNAs and Their Regulatory Roles in Plant– Environment Interactions. Annual review of plant biology 70, 489–525.

Stavolone, L., Kononova, M., Pauli, S., Ragozzino, A., de Haan, P., Milligan, S., Lawton, K., and Hohn, T. (2003). Cestrum yellow leaf curling virus (CmYLCV) promoter: a new strong constitutive promoter for heterologous gene expression in a wide variety of crops. Plant molecular biology 53, 703–713.

Taochy, C., Gursanscky, N.R., Cao, J., Fletcher, S.J., Dressel, U., Mitter, N., Tucker, M.R., Koltunow, A.M.G., Bowman, J.L., Vaucheret, H., and Carroll, B.J. (2017). A Genetic Screen for Impaired Systemic RNAi Highlights the Crucial Role of DICER-LIKE 2 Plant physiology 175, 1424–1437.

To□r, M., Gordon, P., Cuzick, A., Eulgem, T., Sinapidou, E., Mert-Tu□rk, F., Can, C., Dangl, J.L., and Holub, E.B. (2002). Arabidopsis SGT1b Is Required for Defense Signaling Conferred by Several Downy Mildew Resistance Genes. The Plant cell 14, 993–1003.

Vigh, M.L., Thieffry, A., Arribas-Hernández, L., and Brodersen, P. (2021). Nuclear and cytoplasmic RNA exosomes and PELOTA1 prevent miRNA-induced secondary siRNA production in Arabidopsis. bioRxiv,2021.2005.2031.446391.

Wang, X.B., Jovel, J., Udomporn, P., Wang, Y., Wu, Q., Li, W.X., Gasciolli, V., Vaucheret, H., and Ding, S.W. (2011). The 21-nucleotide, but not 22-nucleotide, viral secondary small interfering RNAs direct potent antiviral defense by two cooperative argonautes in Arabidopsis thaliana. The Plant cell 23, 1625–1638.

Wei, X., Ke, H., Wen, A., Gao, B., Shi, J., and Feng, Y. (2021). Structural basis of microRNA processing by Dicer-like 1. Nature Plants 7, 1389–1396.

Wu, Y.-Y., Hou, B.-H., Lee, W.-C., Lu, S.-H., Yang, C.-J., Vaucheret, H., and Chen, H.-M. (2017). DCL2- and RDR6-dependent transitive silencing of SMXL4 and SMXL5 in Arabidopsis dcl4 mutants causes defective phloem transport and carbohydrate over-accumulation. The Plant Journal 90, 1064–1078.

Xie, Z., Allen, E., Wilken, A., and Carrington, J.C. (2005). DICER-LIKE 4 functions in trans-acting small interfering RNA biogenesis and vegetative phase change in Arabidopsis thaliana. Proceedings of the National Academy of Sciences of the United States of America 102, 12984–12989.

Xie, Z., Johansen, L.K., Gustafson, A.M., Kasschau, K.D., Lellis, A.D., Zilberman, D., Jacobsen, S.E., and Carrington, J.C. (2004). Genetic and functional diversification of small RNA pathways in plants. PLoS Biol 2, E104.

Yoshikawa, M., Peragine, A., Park, M.Y., and Poethig, R.S. (2005). A pathway for the biogenesis of trans-acting siRNAs in Arabidopsis. Genes & development.

Yoshikawa, M., Han, Y.-W., Fujii, H., Aizawa, S., Nishino, T., and Ishikawa, M. (2021). Cooperative recruitment of RDR6 by SGS3 and SDE5 during small interfering RNA amplification in Arabidopsis. Proceedings of the National Academy of Sciences 118, e2102885118.

Zhang, X., Zhang, X., Singh, J., Li, D., and Qu, F. (2012). Temperature-Dependent Survival of Turnip Crinkle Virus-Infected Arabidopsis Plants Relies on an RNA Silencing-Based Defense That Requires DCL2, AGO2, and HEN1. Journal of Virology 86, 6847–6854.

Zhang, X., Zhu, Y., Liu, X., Hong, X., Xu, Y., Zhu, P., Shen, Y., Wu, H., Ji, Y., Wen, X., Zhang, C., Zhao, Q., Wang, Y., Lu, J., and Guo, H. (2015). Suppression of endogenous gene silencing by bidirectional cytoplasmic RNA decay in Arabidopsis. Science 348, 120–123.

