## Supplemental Table and Figures for "Evidence for an RNAi-independent role of DICER-LIKE2 in conferring growth inhibition and basal antiviral resistance"

| **Allele** | **Mutagen** | **Alias** | **Aa change** | **Reference** |
|---|---|---|---|---|
| *dcl2-1* | T-DNA | SALK_064627 | | (Xie et al., 2004) |
| *dcl2-2* | T-DNA | SALK_123586.28.70.x | | www.arabidopsis.org |
| *dcl2-3* | T-DNA | SALK_095069 | | www.arabidopsis.org |
| *dcl2-4* | EMS | *rtp5-1* | W796* | (Taochy et al., 2017) |
| *dcl2-5* | EMS | *rtp5*-2 | A1098V | (Taochy *et al.*, 2017) |
| *dcl2-6* | EMS | | P78S | This study |
| *dcl2-7* | EMS | | G106E | This study |
| *dcl2-8* | EMS | | V107M | This study |
| *dcl2-9* | EMS | | G197D | This study |
| *dcl2-10* | EMS | | E665K | This study |
| *dcl2-11* | EMS | | W796* | This study |
| *dcl2-12* | EMS | | G898E | This study |
| *dcl2-13* | EMS | | Q991* | This study |
| *dcl2-14* | EMS | | Q1040* | This study |
| *dcl2-15* | EMS | | G424E | This study |

**Supplemental Table 1** Overview of mutant alleles of the Arabidopsis *DCL2* gene.

**References:**

Taochy, C., Gursanscky, N.R., Cao, J., Fletcher, S.J., Dressel, U., Mitter, N., Tucker, M.R., Koltunow, A.M.G., Bowman, J.L., Vaucheret, H., and Carroll, B.J. (2017). A Genetic Screen for Impaired Systemic RNAi Highlights the Crucial Role of DICER-LIKE 2 Plant Physiology *175*, 1424-1437. 10.1104/pp.17.01181.

Xie, Z., Johansen, L.K., Gustafson, A.M., Kasschau, K.D., Lellis, A.D., Zilberman, D., Jacobsen, S.E., and Carrington, J.C. (2004). Genetic and functional diversification of small RNA pathways in plants. PLoS Biol 2, E104.

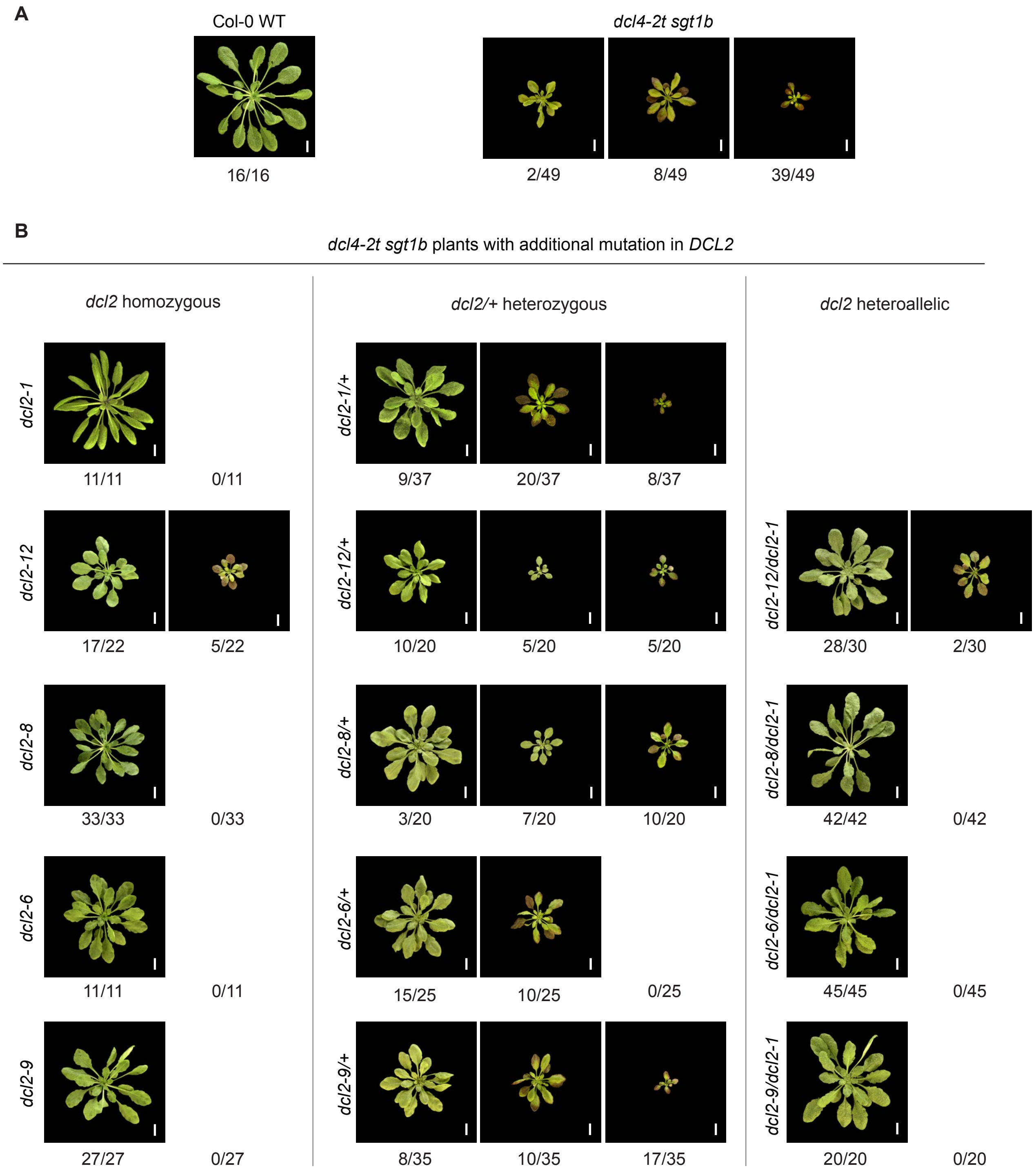


**Supplemental Figure 1. Suppression of *dcl4-2 sgt1b* phenotypes by DCL2 point mutants allowing siRNA production**
**A**, Rosette phenotypes of 49-day old plants of the indicated genotypes grown in short day conditions. Three different categories of *dcl4-2t sgt1b* phenotypes are shown. Scale bars, 1 cm. **B**, Rosette phenotypes of *dcl4-2t sgt1b dcl2-x* plants with the *dcl2* mutant alleles either in the homozygosity (left column), heterozygosity (middle column) or in the heteroallelic *dcl2-1/dcl2-x* combination (right column). *dcl2* heterozygous plants were obtained as F1 progeny of crosses of the reference *dcl4-2t sgt1b dcl2-1* or the newly isolated *dcl4-2t sgt1b dcl2-x* to *dcl4-2t sgt1b*. The heteroallelic combinations *dcl4-2t sgt1b dcl2-1/dcl2-x* were obtained as F1 progeny of crosses of *dcl4-2t sgt1b dcl2-1* to *dcl4-2t sgt1b dcl2-x*. *dcl2* allele numbers are described in Figure 5 and Table S1. Scale bars, 1 cm.

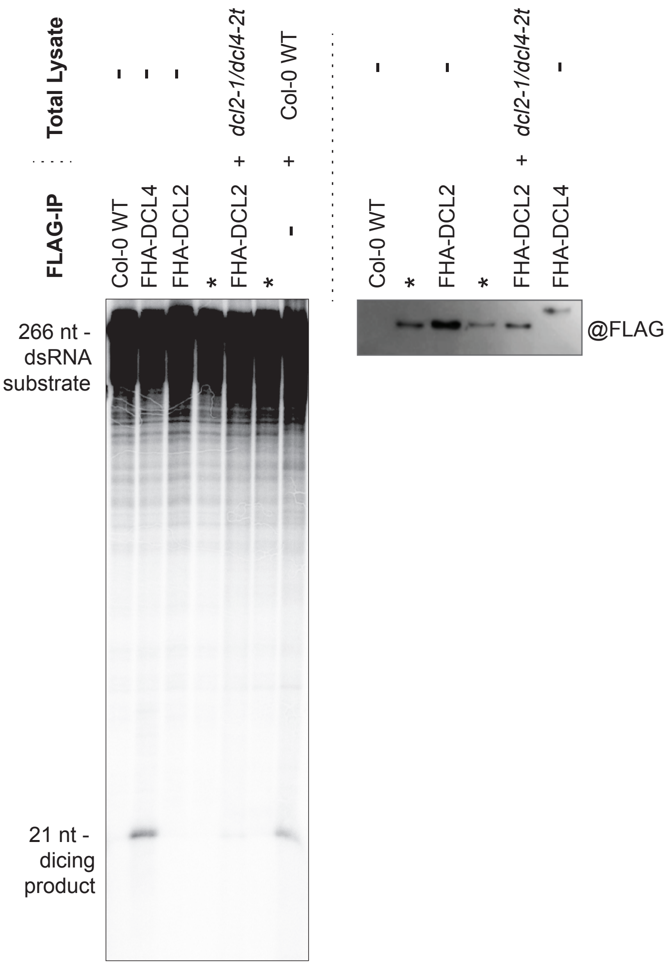


**Supplemental Figure 2.** ***In vitro*** **assays of dicer activity of immuno-purified DCL4 and DCL2**

Dicer activity against long, blunt dsRNA of FLAG-HA-DCL4 (FHA-DCL4) and FLAG-HA-DCL2 (FHA-DCL2) immunopurified from flower lysates. Left panel, autoradiogram showing the separation of cleavage products obtained by incubation of *in vitro* transcribed, radiolabeled dsRNA (perfectly complementary PHB fragments allowed to hybridize to form dsRNA after *in vitro* transcription), with FLAG-immunopurified fractions (FLAG-IP) from total lysates of either non-transgenic plants (Col-0 WT), plants expressing FHA-DCL4 or plants expressing FHA-DCL2. 10 μl of total lysate from *dcl4-2t dcl2-1* plants was added to the indicated enzymatic reactions to provide additional components that DCL2 may need for activity. A reaction containing only Col-0 WT lysate (with endogenous DCLs) was added as positive control (right-most lane). Right panel, western blot developed with FLAG antibodies of 1/3 of the immunopurified fractions used for the Dicer activity assay. Asterisks indicate reactions with FHA-DCL2 immunopurified from transgenic lines not described in this article, and can be ignored for the purpose of this study.
